# Deletion of beta-fructofuranosidase (invertase) genes is associated with sucrose content in Date Palm fruit

**DOI:** 10.1101/652289

**Authors:** Joel A. Malek, Sweety Mathew, Lisa Mathew, Shameem Younuskunju, Yasmin A. Mohamoud, Karsten Suhre

## Abstract

The fruit of date palm trees are an important part of the diet for a large portion of the Middle East and North Africa. The fruit is consumed both fresh and dry and can be stored dry for extended periods of time. Date fruits vary significantly across hundreds of cultivars identified in the main regions of cultivation. Most dried date fruit are low in sucrose but high in glucose and fructose. However, high sucrose content is a distinctive feature of some date fruit and affects flavor as well as texture and water retention. To identify the genes controlling high sucrose content we analyzed date fruit metabolomics for association with genotype data from 121 date fruits. We found significant association of dried date sucrose content and a genomic region that contains 3 tandem copies of the beta-fructofuranosidase (invertase) gene in the reference Khalas genome, a low sucrose fruit. High sucrose cultivars including the popular Deglet Noor had a homozygous deletion of two of the 3 copies of the invertase gene. We show the deletion allele is derived when compared to the ancestral allele that retains the all copies of the gene in 3 other species of *Phoenix*. The fact that 2 of the 3 tandem invertase copies are associated with dry fruit sucrose content will assist in better understanding the distinct roles of multiple date palm invertases in plant physiology. Identification of the recessive alleles associated with end-point sucrose content in date fruit may be used in selective breeding in the future.

## Introduction

The date palm (*Phoenix dactylifera*) is a tree of agricultural significance in a large area stretching from North Africa, through the Middle East and to parts of South Asia. Hundreds of different cultivars are popular in this region, each with distinct taste, color, texture, and nutritional profiles. Dates have provided a significant energy source that is easy to dry, transport and store for people living in the region for millennium. The sugar content of dates has been studied and noted that glucose, fructose and sucrose are the main sugars with their balance changing depending on fruit ripening stage or cultivar type (Cook and Furr 1952; Hasegawa and Smolensky 1970). Dates are categorized by texture into “soft”, “semi-dry” and “dry” cultivars depending on water retention in dry fruit. Others have shown that these texture categories may also relate to the balance of sugars in the dates with higher sucrose associated with the low water retention found in the dry and semi-dry cultivars(Mustafa et al. 1986; Yahia and Kader 2011; Diboun et al. 2015). High sucrose content in dates has been associated with low invertase enzyme levels which are necessary to convert reducing sugars to sucrose(Hasegawa and Smolensky 1970). The date palm genome contains multiple invertases (Al-Dous et al. 2011; Al-Mssallem et al. 2013) for conversion of sucrose in various cellular and physiological pathways, however specifically which invertases are involved in dry fruit sucrose content has not been identified. There is a reduction in date palm genome sequence diversity specifically in regions containing sugar metabolism genes (Al-Mssallem et al. 2013; Hazzouri et al. 2015) possibly indicating selection during cultivation. It would be of interest to identify potential genetic control of sucrose content in dried dates to improve breeding programs and development of post-harvest date fruit quality.

We previously reported the metabolomics analysis of 123 date fruits (Stephan et al. 2018) and their respective genotyping(Thareja et al. 2018). Here we combine these data sets and analyze the sucrose measurements for association with genetic variants for a total of 121 dates. To understand the possible genetic basis of sucrose content in date fruit we conducted an association study between the genome-wide genotypes of 121 date cultivars with their sucrose content at the dried fruit stage.

## Methods

### Sample collection and sequencing

Date fruit and leaves from Phoenix species samples [Supplementary Table 1] were collected from across the date palm growing world, DNA extracted and sequenced as described (Mathew et al. 2014, 2015). Briefly, fruit samples were prepared with the Qiagen Plant DNeasy DNA extraction kit, libraries prepared and sequenced on either the HiSeq2500 or 4000 system according to the manufacturers recommended protocol.

### Sequence analysis

The genomes of the date fruit have been sequenced and variable regions genotyped (Thareja et al. 2018) using BOWTIE2 for sequence alignment and SAMTOOLS for single nucleotide polymorphism (SNP) calling against the date palm PDK30 reference genome(Al-Dous et al. 2011). Indels were not considered and only a single SNP from close clusters of SNPs were selected for analysis resulting in a total of 1.4M SNPs. Annotation of single-molecule sequencing based scaffolds was conducted with MAKER (Cantarel et al. 2008).

### Genotype association with sucrose

Metabolite information from the date fruit samples was collected by Metabolon as previously described (Stephan et al. 2018). The genotype data was tested for association with the date fruit sucrose content (semi-quantitative non-targeted metabolomics data, run-day normalized, and normalized by Bradford protein content) using linear regression as implemented in the PLINK software (version 1.9). We found associations (p<10^−10^) with sucrose on 34 contigs, the strongest association at p=5×10^−16^ and a genomics inflation of lambda=1.28. These contigs were matched to the oil palm reference using MUMmer (Marçais et al. 2018). The highest associated SNPs were located in the 60kb contig PDK30s742521 (http://qatar-weill.cornell.edu/research/research-highlights/date-palm-research-program/date-palm-draft-sequence) which was matched to longer contigs from a single-molecule based date palm reference sequence of both the Khalas (PDKtg42s0452_505576) and Deglet Nour genomes (PDD10s0018_1259392) (Torres et al. 2018). The Khalas reference sequence was noted to include sequence missing in the Deglet Nour reference and so all genomes were aligned to the Khalas contig and diploid coverage normalized to 1 for analysis of copy number changes in the region. Sample relatedness was calculated in VCFTOOLS (Danecek et al. 2011) using the built-in kinship algorithm.

## Results

Association of the date fruit genotype data to sucrose for the 121 date fruit samples (Table 2, Supplementary Table 1 and 2) revealed a region of date palm contigs with similarity to one main region of approximately 1Mb in the more contiguous oil palm genome reference (Figure 1a,b). Investigation of the genes in contigs with the most highly associated SNPs to sucrose content showed that PDK30s742521_60107 contained a predicted invertase gene (Supplementary Table 3) and that annotation of the extended corresponding single-molecule based contig (PDKtg42s0452_505576) showed 3 tandem copies of the invertase gene in the first 100kb (Figure 1c). All high sucrose cultivars were homozygous for the alternative allele to the khalas reference (Figure 2). A contig with the homologous region from the high-sucrose cultivar Deglet Nour was identified (PDD10s0018_1259392) and aligned to the Khalas contig. Deglet Nour was found to be missing a region spanning 2 of the 3 copies of the beta-fructofuranosidase gene (invertase) (Figure 3). To verify that this was not unique to Deglet Nour the top 22 sucrose containing date palm cultivars were selected for further sequence coverage analysis. Upon matching to the Khalas (low sucrose) reference sequence, those cultivars with high sucrose were found to contain no sequences matching the same region deleted in Deglet Nour (Figure 1c,d). As expected, heterozygotes for the deletion contained approximately half the coverage of homozgyotes for the reference allele (Figure 1d). Genotypes of SNPs identified in PDK30s742521_60107 corresponded to the genotype state of the deletion (Figure 4). While multiple high-sucrose samples are simply Deglet Nour collected from different locations, Sukkary (Saudi Arabia) and Naboot Ali were genetically distinct (Supplementary Table 2) with a kinship score of less than 0.25 among these cultivars (Supplementary Table 4). The high-sucrose phenotype appears to be recessive as homozygotes for the deletion were the only group with very high sucrose. Heterozygotes appear to maintain enough invertase activity to convert the sucrose to glucose and fructose (Figure 2).

**Table 1.** Occurrence of the deleted allele by geographical region. Del – homozygous deletion, Het – heterozygous for the reference allele, Hmz – heterozygous for the deletion.

| Location | Genotype | Count | % |
| --- | --- | --- | --- |
| East | Del | 3 | 5.5 |
|  | Het | 1 | 1.9 |
|  | Hmz | 50 | 92.6 |
|  | Total | 54 |  |
| West | Del | 3 | 4.5 |
|  | Het | 15 | 22.4 |
|  | Hmz | 49 | 73.1 |
|  | Total | 67 |  |

**Table 2.** SNPs with high association to the sucrose phenotype. Based on alignment to PDK30 genome. Ref/Alt determination is based on the Khalas reference sequence.

|  |  |  |  |
| --- | --- | --- | --- |
| PDK30s742521_60107 | 39241 | C | G |
| PDK30s742521_60107 | 39533 | A | T |
| PDK30s742521_60107 | 39691 | C | A |
| PDK30s742521_60107 | 40462 | A | C |
| PDK30s742521_60107 | 40498 | T | C,A |
| PDK30s6550997_92357 | 13026 | C | A |
| PDK30s6550997_92357 | 7860 | C | G |
| PDK30s6550997_92357 | 6838 | C | T |
| PDK30s6550997_92357 | 9130 | A | G |
| PDK30s6550997_92357 | 9214 | G | A |
| PDK30s6550997_92357 | 9215 | C | A,T |
| PDK30s6550997_92357 | 9216 | A | C |

**Figure 1.**
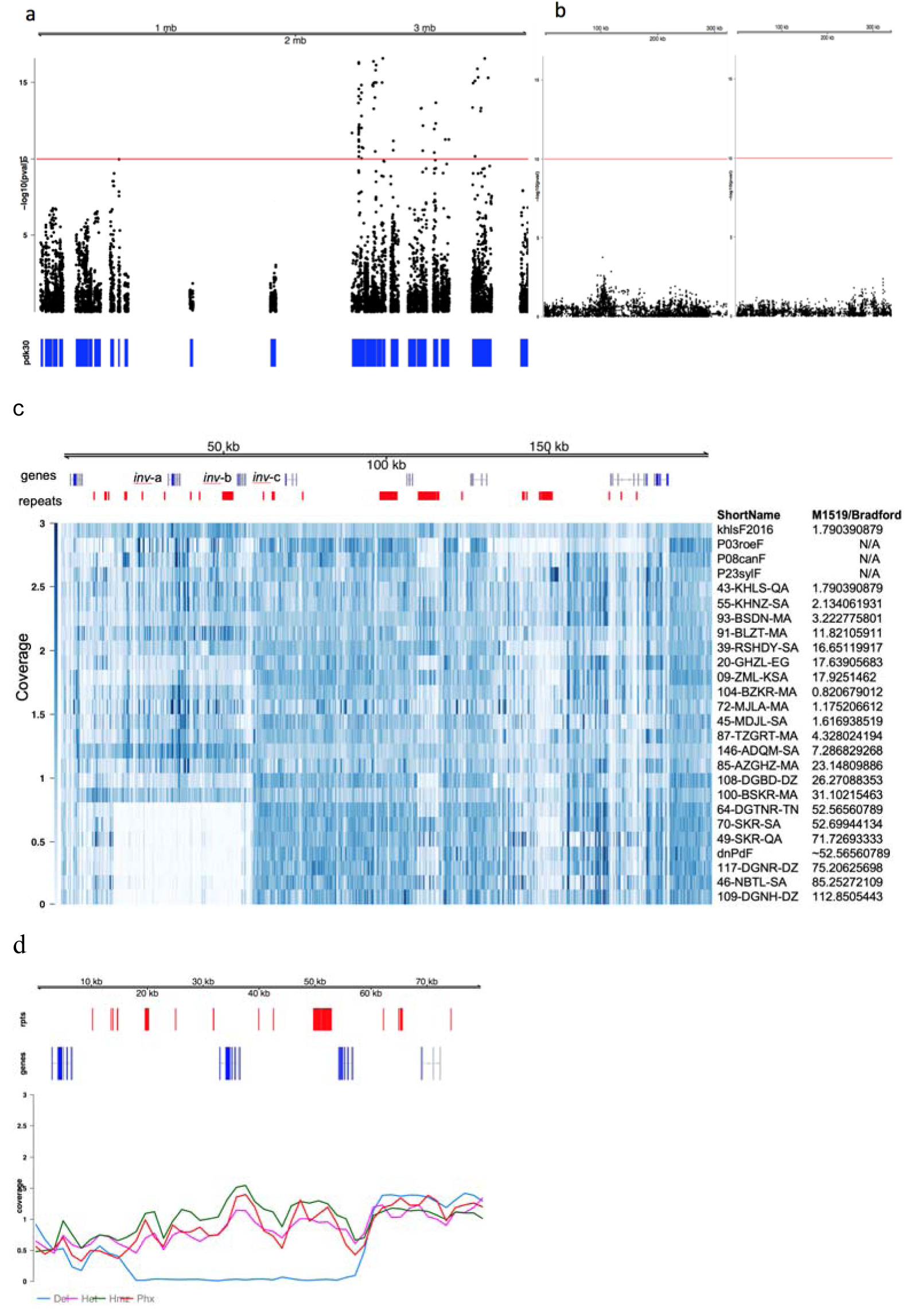
Identification of an allele associated with sucrose content in date fruit. A. Date palm sequence contigs containing SNPs associated with fruit content were aligned to the oil palm reference sequence and found to derive mainly from a 1Mb region of contig NC_026001.1 of Elaeis guineensis chromosome 9. PDK30: Date Palm contigs are represented as blue rectangles. A red line is drawn at the multiple testing corrected genome-wide significance of [-log(10) of 10]. b. Two randomly selected PDK30 contigs with SNPs plotted as in (a) showing SNP p-values do not reach genome-wide significance. c A heatmap of normalized coverage for selected samples from varying genotypes. Samples with high sucrose are shown to be deleted in the region containing the invertase gene. dnPdF is predicted to have this value of sucrose given other Deglet Noor genomes. d. Coverage grouped by genotype in the region of the invertase gene. Lines are loess-smoothed and diploid coverage is normalized to 1. The deleted samples who know coverage in the region while heterozygotes show reduced coverage. Phoenix species are likely homozygous for the allele containing both copies of the invertase but show slightly lower coverage than homozygous date fruit samples likely due to sequence divergence impacting sequence alignment of Phoenix sequences to the Khalas reference..

**Figure 2.**
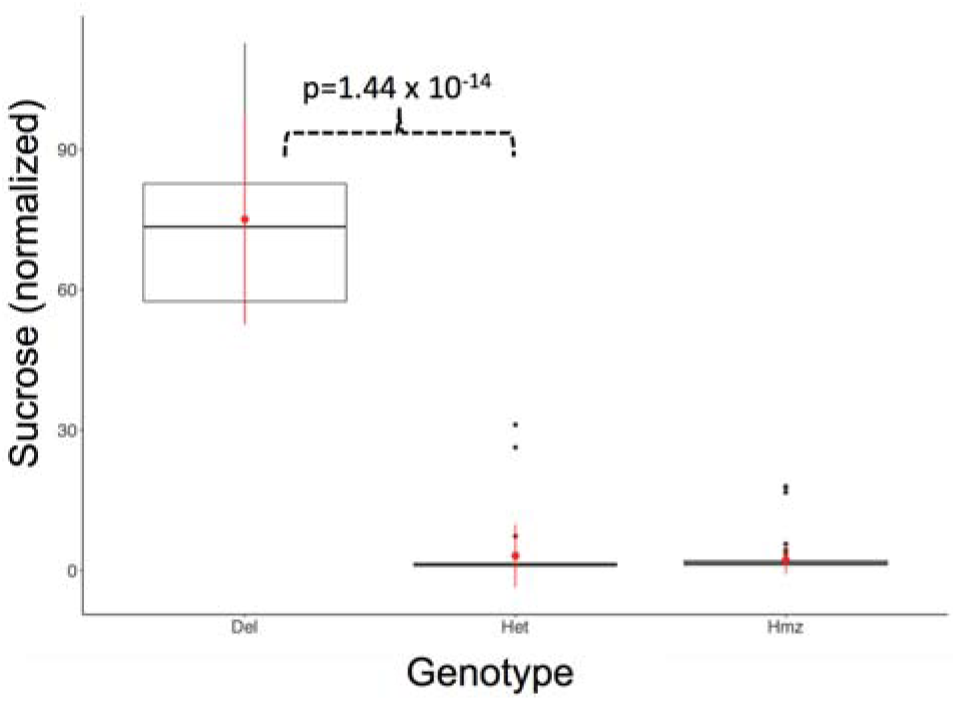
High sucrose content in date palm is associated with a deleted allele. A plot of normalized sucrose content in date fruit for samples deleted for the invertase genes (Del), heterozygous for the deletion (Het) or homozygous for the ancestral, non-deleted allele (Hmz). Genotypes at the position 39533 of contig PDK30s742521_6010739533

**Figure 3.**
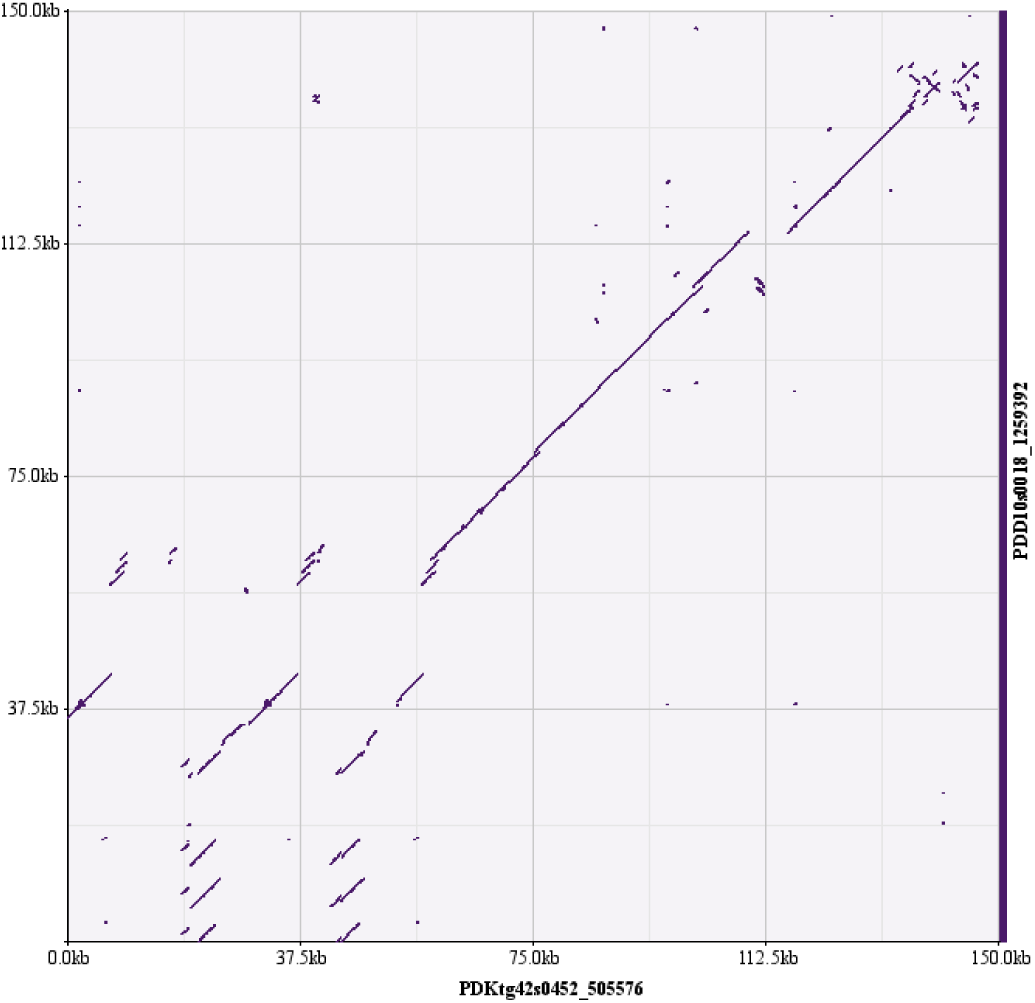
A dot plot of the sequence similarity between the sucrose-associated region in Khalas (low sucrose) and Deglet Noor (high sucrose). Sequences are highly conserved between the two genomes except in the region from 0-50kb where the Deglet Noor genome shows loss of the duplicated invertase genes.

**Figure 4.**
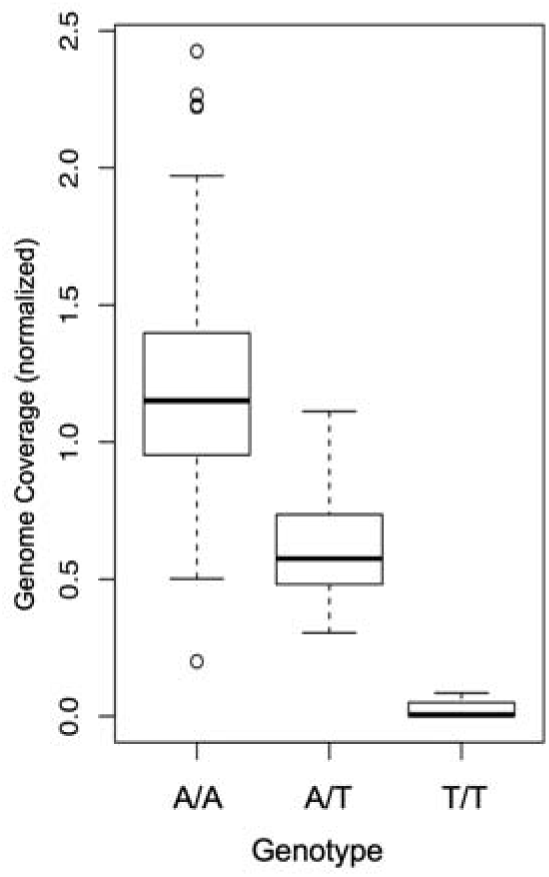
Normalized genome coverage by genotype for PDK30s742521_60107 position 39533

We searched for the presence of the deleted allele in the rest of the date palm population that have been sequenced and found that heterozygotes of the deletion (carriers) are common. The allele is more common in North African cultivars with 18 out of 67 North African cultivars carrying the allele. In contrast, only 4 four out of 54 Eastern cultivars carry the allele (Table 1).

## Discussion

Sucrose and its hydrolyzing enzyme invertase, play critical roles in multiple plant functions including response to the environment and regulation of the life cycle (Sturm 1999; Roitsch and González 2004). Within these functions, vacuolar invertases play a role in the balance of fruit sugar content (Roitsch and González 2004). Research on the effect of invertase in tomato fruit revealed accumulation of sucrose was linked to a recessive invertase mutation (Klann et al. 1993).

The texture of dried date fruit and their classification as soft, semi-dry and dry has been related to sucrose content with low sucrose content dates retaining more water and a softer texture (Cook and Furr 1952; Joseph Kanner et al. 1978; Samarawira 1983; Mustafa et al. 1986). Previous research identified invertase activity as associated with dried date fruit sucrose content (Joseph Kanner et al. 1978) though to-date genetic control of date palm invertase levels specifically in fruit has been elusive. Accumulation of sucrose to high levels in the dried fruit stage has been observed in some cultivars including Deglet Noor. Both Deglet Noor (high sucrose) and Barhee (high glucose/fructose, low sucrose) begin the ripening stage with approximately similar levels of 20% reducing sugars.

Likewise, by the Khala stage of ripening Barhee has 62% dry weight of sucrose and Deglet Noor 40%. However, dramatically, by the fully ripe dried stage (Tamr) Barhee dry weight was only 0.25% sucrose while Deglet Noor’s dry weight was 58% sucrose (Rygg 1946; Samarawira 1983). Date fruit with high invertase activity retain more water through the ripening process, even in dry climates, and this is expected to have an effect on other enzymatic activities affecting fruit features. Our results on sucrose content in dates agrees well with the findings of Zhang and colleagues who found high sucrose in 3 of the same date fruit cultivars including Deglet Noor, Sukkary and Nabtat (Zhang et al. 2015).

Here we show that SNPs associated with very high levels of sucrose content in dried dates are located within a genomic region containing a deletion of two invertase genes. While there are multiple other genes in this region that could potentially play a role in sucrose content, the deletion of two of the invertase genes suggests a central and functional role for these invertase losses in such a strong phenotype. In support of this argument, previous studies on Deglet Noor show that the addition of topical invertase improved fruit texture by removal of crystalline sugars (“sugar wall” dates) that are associated with low invertase levels in the fruit (Smolensky et al. 1975).

This invertase deletion allele appears to be recessive as all high-sucrose content dates contained a homozygous deletion of the invertase genes. This is similar to findings in tomato where a recessive invertase mutation in wild tomato results in high accumulation of sucrose in the fruit (Klann et al. 1993). Interestingly, heterozygotes of the deletion did not seem to have reduced levels of sucrose levels though more research will be required to better understand if this is simply a result of end-point measurement of sucrose content and that the heterozygous deletion does affect earlier ripening stages.

The extremely high accumulation of sucrose in just a few cultivars does not fully explain the link between texture and sucrose content. That is, not all date fruit that are classified as “dry” or “semi-dry” contain high levels of sucrose at the final ripening stages. Observations of sucrose accumulation through the ripening process and not just at the end may play a part in the final texture (Mustafa et al. 1986). Possibly different haplotypes of the 3 invertase genes may be active at different points in time and thereby explain a more complex phenotype. Future association studies on sucrose levels throughout the development process will be important to identify other genes potentially involved in the texture of dates.

In light of recent findings of a strong population division between eastern and western date palm cultivars (Chaluvadi et al. 2014; Mathew et al. 2015; Zehdi-Azouzi et al. 2015; Flowers et al. 2019) it is important to consider which geographical regions this allele is found in. All dates with high sucrose shared the same allele that contained the same structure of deletion and this suggests that this occurred once and spread among the cultivars as opposed to a convergent event between East and West subpopulations. With this in mind, the fact that we observe the allele in samples from both eastern and western subpopulations suggests that enough generations have passed for the allele with invertase deleted to spread in the population and to allow for sufficient genomic mixture such that the Sukkary and Naboot Ali are genetically more eastern cultivars while the Deglet Nour is genetically more western(Chaluvadi et al. 2014; Mathew et al. 2015; Thareja et al. 2018). Where exactly the allele arose is difficult to determine. While we observed it to be more common in North African cultivars, this could simply be a result of multiple cultivars being derived from Deglet Noor. At the same time, despite genomic analysis clearly placing Deglet Noor in the western subpopulation, it has been noted to contain chloroplast DNA from the Eastern subpopulation suggesting Eastern maternal introgression at some point (Zehdi-Azouzi et al. 2015). That it is a derived allele and not ancestral is supported by the fact that we observed related *Phoenix* species to contain all 3 tandem copies of the invertase genes rather than the deletion.

While it is well known that invertase genes play a role in fruit sucrose levels, our findings here help identify the specific genetic basis of dry date fruit extreme sucrose content. The fact that high-sucrose date fruit contains a single copy of the invertase at this location suggests that the retained invertase may be of importance to other functions and pinpoints the final sucrose inversion process to the deleted two genes. Variants in alleles that retain all 3 invertases may play a role in the ultimate texture and water retention of dry dates beyond the extreme phenotype associated with homozygous deletion of the genes. Our findings here could assist future date palm breeding programs and provide the basis for future studies to better understand of the role of specific invertases in the pathway of fruit development.

## Author contributions

JAM designed the study, analyzed data and wrote the manuscript, SM collected samples and analyzed data, LM processed samples and conducted sequencing and analysis, SY conducted DNA sequencing analysis, YAM conducted sequencing and analysis, KS designed the study, conducted metabolomics analysis and wrote the manuscript

## Compliance with Ethical Standards

Human and animal subjects were not included in this study.

The authors declare not competing interests.

## Acknowledgments

This study was supported by grant NPRP-EP X-014-4-001 from the Qatar National Research Fund (a member of Qatar Foundation). We thank Robert Krueger of the USDA, Sean Lahmeyer of Huntington Gardens and Diego Rivera of the University of Murcia for contributing date palm, *Phoenix* and palm species.

